# The importance of semantic network brain regions in integrating prior knowledge with an ongoing dialogue

**DOI:** 10.1101/276683

**Authors:** Petar P. Raykov, James L. Keidel, Jane Oakhill, Chris M. Bird

## Abstract

To understand a dialogue we need to know the specific topics that are being discussed. This enables us to integrate our knowledge of what was said previously, in order to interpret the current dialogue. Here, we selectively manipulated knowledge about the narrative content of dialogues between two people, presented in short videos. The videos were clips taken from television situation comedies and the speech in the first-half of the clip could either be presented normally (high context) or spectrally rotated in order to render it unintelligible (low context). Knowledge of the preceding narrative boosted memory for the following dialogues as well as increased the inter-subject semantic similarity of recalled descriptions of the dialogues. Sharing knowledge of the preceding narrative across participants had two effects on fMRI markers of neural processing: (1) it strengthened temporal inter-subject correlations in regions including the left angular (AG) and inferior frontal gyri (IFG), and (2) it increased spatial inter-subject pattern similarity in the bilateral anterior temporal lobes (ATL). We argue that these brain regions, which are known to be involved in semantic processing, support the activation and integration of prior knowledge, which helps people to better understand and remember dialogues as they unfold.

## Introduction

Understanding a dialogue between two people requires multiple inter-related cognitive processes, yet it is something that we typically accomplish with remarkable ease. In addition to processing the speech, gestures and expressions of the speakers, we need to activate relevant prior knowledge in order to understand what is currently being talked about. This is essential in the situation where a dialogue is resumed after a break; to pick up the conversation, we need to remember the exact topic of the preceding discussion. In this study, we investigate how knowledge of the earlier narrative context affects comprehension and memory for dialogues between people. More generally, we examine the brain regions that integrate prior knowledge from long-term memory with incoming narrative information.

Our study focuses on how we processes observed conversations that are taken from television situation comedies and is related to studies investigating the processing of narratives more generally (see Lee et al., 2020; Willems et al., 2020) The aim is to establish how people are able to comprehend and remember dialogues that are continuations of earlier conversations – using the knowledge gained from the first half to better understand the second half of the dialogue. This is a prerequisite for many day-to-day situations such as continuing an interrupted meeting or when watching episodes from a television show. Nevertheless, it is well-established that actively engaging in a conversation involves collaboration by the speakers to coordinate their beliefs about common ground (Schober & Clark, 1989; Wilkes-Gibbs & Clark, 1992). Therefore, our study provides a starting point for understanding the cognitive and neural processes at play when people converse with each other (see also Stephens et al., 2010).

A dialogue about a specific topic is an example of a single event; the same people are engaged in the same action at a particular time and place (Zacks et al., 2007). These stable elements of a particular situation are described by “event models”, and the event model summarising the current situation – what is happening now – is thought to be held within working memory (a so-called “working event model”; Zacks, 2020). MRI scanning combined with the use of naturalistic narrative materials, such as movies and stories, has shed light on the brain regions that represent event models (see Bird, 2020; Hasson et al., 2015; Lee et al., 2020). It is well established that “long-timescales regions” (Hasson et al., 2015), which overlap substantially with the so-called “default mode network” (the DMN; Raichle et al., 2001), integrate complex narrative information over time periods of tens of seconds and minutes. This is assumed to reflect the continued maintenance and updating of information about the current situation – the working event model. However, relatively few studies have focussed on situations where narrative information must be integrated across delays, when the information is no longer in working memory and has to be retrieved from long-term memory (though see Chang et al., 2021; Cohn-Sheehy et al., 2020; Lahnakoski et al., 2017).

A popular method for investigating the influence of prior information on the comprehension and memory of events, is to use ambiguous or confusing prose passages or movies, and provide background contextual information that enables the participant to understand them (e.g. Bransford & Johnson, 1972; Dooling & Lachman, 1971; Maguire et al., 1999). Ames and colleagues (2015) used such a manipulation and showed that participants sharing the same background knowledge showed increased synchronization in BOLD response in the ventromedial prefrontal cortex (vmPFC) and posterior cingulate cortex (PCC) when listening to vignettes. It is important to note that here and elsewhere, the term “share” refers to situations where participants have been exposed to the same information or interpret narrative material in a similar way. Moreover, “shared” fMRI responses are responses that a significantly correlated across individuals who are engaged in the same activity (see Chen et al., 2016, 2017; Nastase et al., 2019). Conceptually similar studies using movies also found that activity in the vmPFC was synchronised across participants who shared knowledge of the first half of the movie (Chen et al., 2016; van Kesteren et al., 2010). However, the key contrast in such paradigms involves comparing meaningful events with events that are - at least initially - difficult to understand. It is therefore unclear whether the effects reflect this more fundamental difference in the conditions (coherent versus incoherent narratives (see also Heidlmayr, 2020)).

Other studies have addressed this issue using narrative materials that are open to different interpretations. For example, Yeshurun et al., (2017) manipulated the interpretation of an ambiguous narrative by telling participants either that the main character’s wife was cheating on him, or that the main character was paranoid. Participants sharing the same interpretation showed similar brain activity in regions of the DMN (see also Lahnakoski et al., 2014), suggesting that these regions represent aspects of the narrative such as the beliefs and emotional states of the characters in the story. In another study, participants viewed two shapes interacting, and those who interpreted the meaning of the interactions more similarly also showed more tightly synchronised brain activity (Nguyen et al., 2019; see also Saalasti et al., 2019 for a similar result with a prose passage). Taken together, there is strong evidence that when people share a common interpretation of a complex situation, brain activity within regions of the DMN is synchronised across individuals. However, these studies have not addressed the issue of how prior knowledge specifically affects how dialogues are processed.

Keidel and colleagues (2017) more directly examined how prior knowledge is integrated with on-going narrative processing to aid comprehension of dialogues. Participants saw first and second halves of clips taken from TV situational comedies (sitcoms). The second half clips could either be a direct continuation of the first half clips, or be clips taken from the same show involving the same characters and location, but from a different episode of the show (and consequently depicting a different narrative storyline). Provision of narrative context resulted in increased activation in regions of the DMN, particularly those associated with semantic processing (Binder et al., 2009; Lambon Ralph et al., 2016; Noonan et al., 2013), including the inferior frontal gyrus (IFG), middle temporal gyrus (MTG) and angular gyrus (AG). It was argued that this pattern of activation reflected the reactivation of linked semantic concepts that were created when watching the first half videos. However, because the non-continuation clips clearly showed a different event, the activation differences may have reflected other differences between the conditions, such as the appearance of the characters and the contents of the scene. The present study avoids these potential confounds by using a modified, but similar, paradigm.

Most of the previous studies identify the brain regions involved in processing complex naturalistic situations by measuring inter-subject correlations (ISCs) in BOLD activity. High ISCs occur when participants either share the same prior knowledge or the same interpretation of an event. More recently, it has been shown that participants often show correlated spatial patterns of brain activity when viewing or listening to the same narrative (inter-subject pattern similarity, ISPS; Chen et al., 2017; Koch et al., 2020; Oedekoven et al., 2017; Wang et al., 2020). Similar effects have been seen when people construct an event in their mind’s eye while listening to it being described (Zadbood et al., 2017). Unlike ISCs, ISPS is based on the multivoxel pattern of brain activity averaged across a whole event lasting 20 – 100 seconds. It is therefore possible that these two inter-subject measures might be sensitive to differences in content of narrative materials. ISCs are driven by temporal variation in activity, and therefore might be driven by particular events within more extended episodes. By contrast, ISPS might reflect aspects of the event models that are constant across the whole episode (see Chen et al., 2017).

The present study uses a similar design to Keidel et al., (2017). We presented participants with video excerpts taken from sitcoms that were divided into first and second halves, in two different conditions: High vs. low context. In the high context condition, the first-half clips were presented with normal speech, whereas in the low context condition we spectrally rotated the speech to render it unintelligible. Under this design, all the pairs of clips were clearly taken from the same episode and differed only in the provision of knowledge about the content of the dialogue. Importantly, all second-half clips, which were the focus of the analyses, were identical for both conditions, with the only difference being the background narrative knowledge that the participants already have before watching the videos. We hypothesized that provision of relevant prior narrative knowledge would aid the comprehension and memory of the dialogues. Indeed, it is possible that the lack of prior narrative knowledge in the low context condition could result in some confusion and misinterpretations when watching the second half video clips. Second, we predicted that participants who shared the same knowledge about the dialogues would activate this knowledge in a more consistent manner, resulting in increased ISCs and ISPS, particularly within regions associated with semantic processing (Binder et al., 2009; Fairhall & Caramazza, 2013; Keidel et al., 2017; Lambon Ralph et al., 2016; Noonan et al., 2013).

## Methods

### Participants

Twenty-four right-handed native English speakers with normal or corrected to normal vision were included in the study. We recruited 24 participants based on prior work (Ames et al., 2015; Chen et al., 2017; Heidlmayr et al., 2020; Keidel et al., 2017; Saalasti et al., 2019; van Kesteren et al., 2010) using ISC and ISPS and work comparing the effects of prior knowledge on processing of naturalistic stimuli. It has been suggested that at least 4 participants per condition are needed to estimate reliable ISC (Hasson et al., 2004). Four participants were not included in the final analysis due to artefacts in the MRI scans and 1 further participant did not complete the experiment. One participant had corrupted post-scanning behavioural data for the memory questions and was not included in the behavioural analysis. The project was approved by the Brighton and Sussex Medical School Research Governance and Ethics Committee and all participants gave informed consent and were paid £20. Additionally, we collected data for a follow-up behavioural study from a separate group of participants that completed the task online. Specifically, we recruited 207 (42 Males, 165 females) participants who were undergraduate psychology students at the University of Sussex. Participants had a mean age of 19.61 (± 2.0). We excluded 38 participants from the study because they failed to recollect more than 30% of the videos used in the study. Therefore 168 participants were included in the main analysis (the overall findings are consistent whether or not these 38 participants are included in the analyses). Participants received course credits in exchange for their participation. The project was approved by the University of Sussex Cross School Research Ethics Committee.

### Stimuli

Twenty video clips from different US and UK television shows were used in the experiment. The TV shows originally aired between 1970 and 2003 and were selected to be unfamiliar to our sample. Each video was divided into first and second halves. The scene location and characters remained constant across the two halves. For our main experimental manipulation, the speech in 10 of the first half videos was made unintelligible. This was done with Praat (version 6.0.15) (Boersma, 2001) by spectrally rotating the audio of the videos with a sinusoidal function with maximum frequency of 4 KHz. This keeps the intonation and rhythm of the speech but makes it incomprehensible. The audio for all videos was scaled to have the same mean decibel intensity. The mean duration of all the excerpts was 32.47(±3.88) seconds. The first half videos (30.57±4.38) were on average shorter than the second half videos (34.37±2.02). Three different video clips, all with spectrally rotated audio, were used in the practice task with total video length of 4.36 minutes.

### Procedure

#### FMRI Study

Participants were informed they would see first- and second-half videos in four separate runs (five full videos in each). They were told that the speech in some of the first-half videos would be unintelligible and were asked to watch the clips as they would watch television at home. Participants were also informed that their memory for the clips would be tested after the scanning session. They completed an 8-minute practice session outside the scanner, to familiarise themselves with the task and the sound of spectrally rotated speech.

Four lists were created to fully counterbalance the conditions and presentation order. For instance, a first half video was presented with normal comprehensible speech (NS see Fig. 1) for half of the participants and the same video was presented with spectrally rotated speech (SRS see Fig.1) for the other half participants. This meant half of the participants had knowledge of the narrative theme for a second half clip (HC see Fig. 1) and the other half did not (LC see Fig. 1). Notably, watching the first half clip even with spectrally rotated speech still provided some information about the event. Thus, even though participants could not understand the dialogue between the actors, the locations, people present and the emotional tone of the event were still obvious in the first-half videos with spectrally rotated speech (e.g. participants would be able to see that a video depicted two smartly dressed men having a convivial conversation at a table in a café, despite the conversation itself being unintelligible). Therefore, all participants had prior knowledge of the gist of the situations when watching the second half clips: the key difference between the HC and LC conditions was the availability of the prior narrative knowledge (see for example stimuli https://tinyurl.com/3b42fzzd). As a consequence of this, participants watching videos in the LC condition may find them more confusing and/or misinterpret aspects of the dialogue. Apart from counterbalancing the conditions which created two lists we also counterbalanced the order in which the videos were presented both across runs and within run. This created two more lists leading to 4 counterbalancing lists.

**Figure 1.**
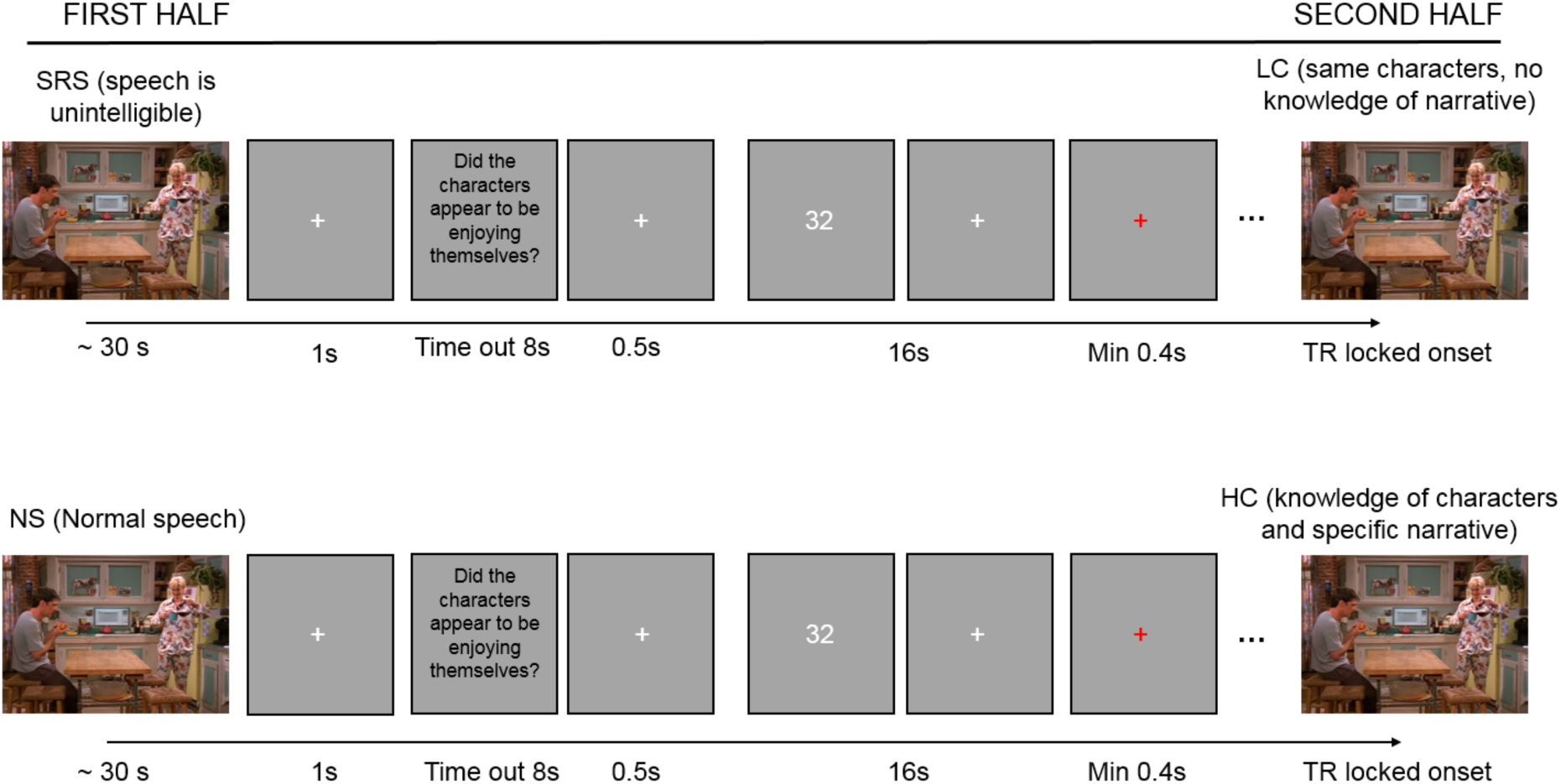
Schematic of study design. Participants viewed videos from unfamiliar TV sitcoms that were divided into two halves. Participants viewed a set of 5 first half videos followed by a set of 5 corresponding second half videos presented in random order. Ten of the first halves had comprehensible speech, whereas the other 10 had unintelligible speech, created by spectrally rotating the audio. Videos were counterbalanced across participants in a within-subjects design. SRS – spectrally rotated speech; NS – normal speech; HC – high context; LC – low context.

The scanning session started with a 4-minute resting state scan (data not reported here). Following this, the task was presented in four functional runs, each approximately 10 minutes long. Each run consisted of 5 first half and 5 second half clips and every clip were followed by a question and an active baseline task. Participants initially watched the set of 5 first half clips and afterwards watched the set of corresponding second half clips in random order. This meant that a second half clip did not immediately follow its corresponding first-half clip. In each run there were either two or three clips with rotated speech. Movie onsets were time-locked to the repetition time (TR). After each clip, a white fixation cross was presented for 1 second. Each video was followed by a question about the general relationship between the characters in the scene (e.g., Did the characters appear to be enjoying themselves?). Behavioural piloting confirmed that participants could answer the questions regardless of whether speech was rotated or not. The questions were presented for 8 seconds or until participants made a Yes/No response and were followed by a 500-millisecond fixation. To obtain an activity baseline, the participants were required to make odd/even judgements about a series of numbers (Stark & Squire, 2001). The active baseline task was presented in 16-second blocks between clips. Six randomly chosen numbers, between 1 and 98, were sequentially presented. Each number was onscreen for 2 seconds and was followed by a fixation cross presented for 667 milliseconds. The active baseline task was used to prevent participants from rehearsing the information presented in the clips. After the baseline task a red fixation cross was presented for a minimum of 400 milliseconds to signal that the next video was about to begin.

Outside of the scanner, participants completed a questionnaire about their familiarity with the shows. Only 6 participants reported any familiarity with 1-3 of the 20 shows. This represented only 3.9% from the data used in the analysis. None of the participants were familiar with the particular scenes used in the experiment, which was our main interest as the first half clips provided some familiarity with the social relationships between characters in the second half clips. Afterwards, participants carried out a computer-based memory task, in which the first 4-6 seconds of the second half videos were presented as a memory cue. Participants were then asked to rate from 1 to 10: (1) the vividness of their memory of the video, (2) how coherent they found the story in the videos, (3) how engaging they found the video. Participants were also asked an open-ended memory question specific to each second half video (e.g., What was the address on his chest written in?). These questions could all be answered from seeing the second half clips alone, but we hypothesised that knowledge of the preceding dialogue would nevertheless boost performance. Responses were rated as either correct or incorrect. All memory questions concerned only material presented in the second half of the videos.

#### Online study

To test in more detail the content that could be recalled from the second half clips from the HC and LC conditions, we additionally ran an online experiment using a very similar design to the fMRI task. The only substantial change was the additional of free recall tests at the end of the experiment. Participants once again viewed first- and second-half video clips in blocks of five (five first half followed by five second half clips). They were informed that some of the first half clips would be with unintelligible speech and were asked to pay attention to all video clips as their memory for the videos would later be tested. An odd-even number task was again included between the clips.

Following the encoding of the videos, participants were shown a screen indicating that the recall session would begin. Memory for videos was tested in a randomized order. Participants were presented with the first 4-6 seconds of the second half videos as a memory cue. Participants were afterwards asked to provide subjective ratings on a scale of 0-100: (1) How vividly they remember the video, (2) How coherent did they find the second half video clip. Then, participants were asked to type in as much detail as possible what they remember from the second half video clip. This was followed by the same open-ended questions used the main experiment.

### MRI acquisition

Data were acquired on a 1.5 T Siemens Avanto MRI scanner. Functional T2* weighted BOLD-sensitive images were acquired with EPI sequence with the following parameters: FOV = 192mm, TR = 2.62 seconds, TE = 42 milliseconds, 90 degree flip angle, slice thickness = 3mm, 35 interleaved ascending slices with .6 mm gap, and 3.0×3.0×3.0 mm voxels. A high resolution T1-weighted image was acquired with the following parameters: FOV 256mm, TR = 2.73 seconds, TE = 3.57ms, 1.0×1.0×1.0mm voxel size.

### Image pre-processing

All EPI images were pre-processed using SPM 12 (Wellcome Department of Imaging Neuroscience, London, UK). Field maps were used to correct for image distortions and susceptibility-by-movement effects using the Realign and Unwarp option (Andersson et al., 2001; Hutton et al., 2002). All EPI images were aligned to the first image of the first session. The anatomical image of each subject was co-registered to their mean realigned EPI images. The anatomical images were then segmented into grey and white matter maps. Anatomical and EPI images were normalized to the MNI space using DARTEL (Ashburner, 2007) and smoothed with an 8 mm FWHM kernel. Images for the inter-subject pattern similarity were pre-processed as above with the exception that a 6 mm FWHM smoothing kernel was applied to the normalised images, as previously used by Chen et al. (2017).

### Data Analysis

Behavioural data was analysed using R and mixed effect models were fitted using the lmer and lmerTest packages. Data were analysed with SPM 12, the CoSMoMVPA toolbox (Oosterhof et al., 2016) and custom scripts in MATLAB (Version 2016b, The MathWorks, Inc., Natick, MA, USA). Permutations tests for whole brain analyses were conducted with command-line functions in FSL (Nichols & Holmes, 2002; Winkler et al., 2014). All analyses were conducted on MNI normalised images within a grey matter mask. Segmentation of the high-resolution structural images provided us with grey matter tissue probability map for each subject. These probability maps were normalised to MNI, averaged across participants. The averaged mask was smoothed with an 8mm FWHM kernel. We selected all voxels within this average probability map higher than a threshold of 0.3 (Nastase et al., 2019). To describe and visualise our data we used the Bspmview toolbox (www.bobspunt.com/bspmview), which implements the MNI coordinates from the Anatomical Automatic Labelling 2 toolbox for SPM 12. Significance was tested with a one-sample random effects t test against zero. Unless otherwise stated, images were whole brain cluster corrected for FWE p < 0.05 at voxel height-defining threshold of p < 0.001.

#### Behavioural Data analysis

Participants who completed the fMRI study had their memory tested outside the scanner. For each second half video clip participants had to answer an open-ended detailed memory question specific to the video (e.g., What was the address on his chest written in?). The second half video clip contained the information necessary to answer the question correctly. For each video we rated whether the participant had answered the question correctly or not. We then submitted the trial level answers to these memory questions to a logistic mixed effect model including random intercept effects for subjects and videos and a random slope effect for subjects and videos: Memory_Question ∼ Condition + (Condition | Subject) + (Condition | Video). The condition indicated whether the video was seen in an HC condition or an LC condition. Separate linear mixed effect models with a fixed effect for the condition and random intercepts for subject and video were fitted for the coherence, vividness, and engagement ratings, due to convergence issues.

In the online study, participants completed a free recall memory test for each of the second half video clips. We rated each trial as either remembered or not. Trials on which participants did not provide any correct details of the video or did not provide information that was not present in the memory cue video were rated as not-remembered and given a rating of 0. Trials on which participants correctly remembered the gist of the clip and provided some details about the clip were provided with a remembered rating (rating of 1; e.g. “The women in the clip claims that penguins cannot fly because their wings cannot support their body-mass. The man disagrees and claims that the sky used to be filled with penguins.”).

For the free-recall data we fitted a mixed-effects logistic regression predicting the free recall data from the condition indicator variable as fixed effect and including random intercept effects for both the subject and videos and included a random-slope effect across participants and videos: Recall ∼ Condition + (Condition | Subject) + (Condition | Video). Linear mixed effect models with the same predictors were fitted for the vividness and coherence data, respectively.

Importantly, the free recall data allowed us to examine whether participants provided more consistent descriptions for the HC videos when compared to the LC videos. Specifically, we used recently developed Google’s Universal Sentence Encoder (USE) to examine semantic similarity for remembered trials in the HC and LC conditions (Cer et al., 2018). First, we selected only remembered trials for the HC and LC conditions. We then converted the memory responses of participants into vectors of 512 numbers using Google’s USE (Cer et al., 2018). The USE allows for whole sentences to be represented as large vectors of numbers. Specifically, these sentence specific vectors are constructed in such a way that the semantic similarity between sentences in maintained and resembles similarity judgments given by people. Such numerical embeddings of text are useful because they allow to quantify similarity of text responses.

After embedding each recollected trial into this semantic space, for each video we extracted the recall responses that were given by participants that saw the video in the HC condition and the other participants that saw the same video in the LC condition. For example, for video one, 74 participants correctly remembered it and had seen it as HC video, and another 71 participants correctly remembered it and had seen it as LC video. For each video we created two average semantic vectors, one for the HC condition and one for the LC condition. We then computed the similarity between each participant’s embedded recall description to the average description of the same video seen in the same condition. We used Person’s correlation coefficient to compute similarity between the embedded recall descriptions. Note the average descriptions were always recalculated to not include the description of the current participant (using a “leave one subject out” procedure). For instance, for video one, we would compute the similarity between the response of participant 1 for that video and the average responses of all *other* participants that remembered that video in the same condition. We then Fisher transformed and averaged these similarity scores across participants to result in two average consistency scores per video (one for the HC condition and one for the LC condition). We than compared for each video whether participants on average provided more consistent responses for the HC condition than the LC condition using a non-parametric permutation test.

### Functional MRI data analyses

#### Univariate General Linear Model (GLM) analysis

A single task regressor for each of the four conditions (SRS, NS, LC and HC: see Figure 1) was included in the first-level models. For all GLM first level models the questions after each video were modelled with a single regressor of no interest and the odd/even number judgment task was left unmodelled to represent the implicit baseline. A block design first level analysis was conducted to replicate previous findings. In this analysis, all video stimuli were modelled with boxcar functions whose durations matched the stimulus duration. The models also included the six motion parameters, a regressor for the mean session effects, and a high-pass filter with a cut-off of 1/128 Hz. We also ran an analysis identical to the above but modelled only the onset of the videos with a gamma function, rather than including the whole duration of the video. This was done to replicate previous findings by Keidel et al. (2017). First-level models for the inter-subject pattern analysis included the same nuisance regressors as the previous analysis. However, each video was represented with its own block regressor that covered the whole duration of the video. This allowed us to examine video specific patterns.

#### Inter-Subject Analyses

##### Inter-subject correlation (ISC)

The ISC allowed us to examine how dynamic processing of the second half videos was modulated by previous knowledge. ISCs were computed voxelwise over second half videos (HC and LC videos, which contained coherent speech). To examine the similarity across participants under the same condition we constructed 2 condition lists. To construct 2 condition lists from 4 counterbalancing lists we combined the lists in which videos were seen under the same condition, but in a different order. There were 9 and 10 people in the two condition lists. It has been shown that averaging over at least 4 people’s time courses provides reliable ISC estimation (Hasson et al., 2004). The first 2 TRs (5.24 seconds) of each video were removed in order to remove transient onset effects that can lead to artificially high ISCs (Ames et al., 2015).

The raw time course for each video and each subject was extracted. These time-courses were used to compute the Fisher-transformed correlations across subjects for each video. For a given subject we computed the correlation between the subject’s specific video time-course and the average time-course for the rest of the participants watching the same video in the same condition (i.e. the participants in the same condition list). This resulted in 20 (10 HC and 10 LC videos) time-course correlations for each subject, which represented the time-course similarity across participants watching the same videos (diagonal values in Fig. 4 A & C).

Apart from ISCs between matching videos we also computed across subject synchronization when participants were watching different videos (e.g. correlating the time-course of a subject watching ‘Dharma and Greg’ with the average time-course of other subjects watching ‘Just Shoot Me’) (see off-diagonal in Fig. 4 A). The ISCs among mis-matching videos acted as a baseline and allowed us to examine ISC while participants watched the same videos irrespective of whether the context was familiar or not (see Fig. 4 A). We compared the time-course correlations across participants watching the same video versus different videos. This general ISC analysis replicates previous work that has examined synchronization across participants experiencing the same event.

For our main analysis, we examined how prior knowledge of the dialogue topics affected ISCs. We examined whether participants were more synchronised when they were watching the HC clips versus when they were watching the LC clips. Each subject had 10 ISCs for the HC clips and 10 ISCs for the LC clips (see Fig. 4 B). These subjects’ values represent his similarity with the rest of the people that watched the same clips under the same conditions. The HC vs LC contrast was performed for each subject by averaging over the 10 Fisher transformed correlations across HC videos and subtracting the average of the 10 LC coefficients. This condition difference was computed for each voxel and for each subject. Therefore 19 subject-specific brain maps were used in the group analysis.

The resulting 19 subject specific brain maps however are not necessarily independent. This is due to the fact that when comparing the correlation between a subject’s time-course and the mean of the “others” there is overlap in the information used to compute the mean “others” across subjects. To illustrate this point: if we have 1-20 subjects within a group, the ISCs for subject 1 is between subject 1 and the average time-course of subjects 2, 3-20. The ISCs for subject 2 involves the average of everyone else, which is 1, 3-20. Therefore, the data for subjects 3-20 were used to calculate the ISCs for both subjects 1 and 2, which means that their ISCs are not independent. Indeed for n subjects in a group, any pair will share n-2 elements (Aly et al., 2018). Because of this, we used non-parametric permutation tests to compute the significance at the group level. The difference in Fisher transformed ISCs between conditions (same vs different videos and HC vs LC) was computed for the general and context specific ISC analyses respectively. To perform the permutations the sign of the resulting difference was flipped for a random subset of subjects before computing the group mean. This effectively is the same as shuffling the conditions for different subjects. 5000 permutations were run (per analysis) to obtain the null distribution with which to compare our observed data and obtain p-values. Cluster corrected images at FWE p < 0.05 at voxel height-defining threshold of p < 0.001 are presented in Figures 4 and 5.

##### Inter-Subject Pattern Similarity (ISPS)

This analysis was performed to examine spatial pattern similarity across participants experiencing the same events. The inter-subject pattern (ISPS) analysis was conducted on the normalised t-maps generated for each subject for each 2nd half video. For each subject, a searchlight map was generated by centring a spherical searchlight with radius of 3 voxels (mean size 110 voxels) at each voxel. For a given participant the activity patterns for a specific video were correlated with the average activity pattern across participants watching the same video in the same condition. This resulted in 20 (10 HC and 10 LC videos) correlations for each subject representing the video specific pattern similarity across subjects (see diagonal Fig. 5 A). The across participants’ pattern similarity was also calculated for non-matching videos resulting in 380 correlation coefficients for each searchlight (see off-diagonal Fig. 5 A). Contrast matrixes were used to weight the resulting Fisher-transformed spatial-pattern correlations for each searchlight. The summed correlations were assigned to the central voxel of each searchlight. Two ISPS analyses were conducted by specifying different contrast matrixes (see Figure 5). Therefore, the central voxel of each searchlight represented the difference between conditions (same vs different videos or HC vs LC) for the general and context specific ISPS respectively. For each of the two analyses we obtained a single condition difference brain image per subject. These brain maps were used in the group analyses.

First, we conducted a “General Show” ISPS analysis to examine the similarity across participants watching the same clip versus watching non-matching clips, irrespective of their prior knowledge (see Fig. 5).

Then we examined whether participants showed more highly correlated patterns for HC videos than for LC videos (see Fig. 5). Due to stimulus counterbalancing, there were two groups (n = 9 and n = 10) of people who saw the same videos in the same condition. Therefore, similarly to the ISC analysis, for the ISPS we compared the similarity between a subject’s patterns and the average patterns of the other subjects in their group who viewed the same video under the same condition.

As above, we again used non-parametric permutation testing to examine the group significant results for the ISPS analyses. This was done by flipping the sign of the ISPS condition difference images for a random subset of subjects. 5000 permutations were run. Results were cluster corrected at FWE p < 0.05 with voxel height defining threshold of p < 0.001.

Both ISCs and ISPSs analyses were also calculated separately for the first half videos using identical procedures to those described above. These analyses examined the synchronization across participants watching the same first half videos. We also contrasted whether participants were more synchronized when they were watching the normal speech (NS) first half videos when compared to videos with spectrally rotated audio (SRS). These analyses showed higher synchronization (both ISC and ISPS) across participants when watching the NS videos compared to the SRS videos in bilateral ATL and other regions often associated with the DMN or semantic processing (see Supplementary Figures 5 and 6).

Based on recent studies examining reinstatement effects in hippocampus when participants were listening to stories referring to previously heard information (Chang et al., 2021; Cohn-Sheehy et al., 2020), we examined whether we would observe higher ISPS effects in hippocampus for HC videos when compared to LC videos. These analyses are reported in supplementary materials.

## Results

### Behavioural results

Participants had an overall accuracy of 72.2% for the memory questions, which is high level of performance given the fact that the questions were open ended (see Fig. 2). Participants responded more accurately to the same questions when in the HC condition (80%) than in the LC condition (64.4%: t_17_ = 3.39, p = 0.003). We fitted a mixed-effect logistic regression including a random intercept and slope for subjects and videos. This analysis suggested that the memory benefit for the HC videos over LC videos would generalise to other stimuli (Z = 2.02, p = 0.04). Due to convergence issues the subjective ratings data were modelled with simple random effect structure including only random intercept for subject and videos. HC videos were also rated as being remembered more vividly (t_323.04_ = 4.42, p < 0.001), more coherent (t_324.86_ = 4.83, p < 0.001) and more engaging (t_323.77_ = 4.62, p < 0.001).

**Figure 2.**
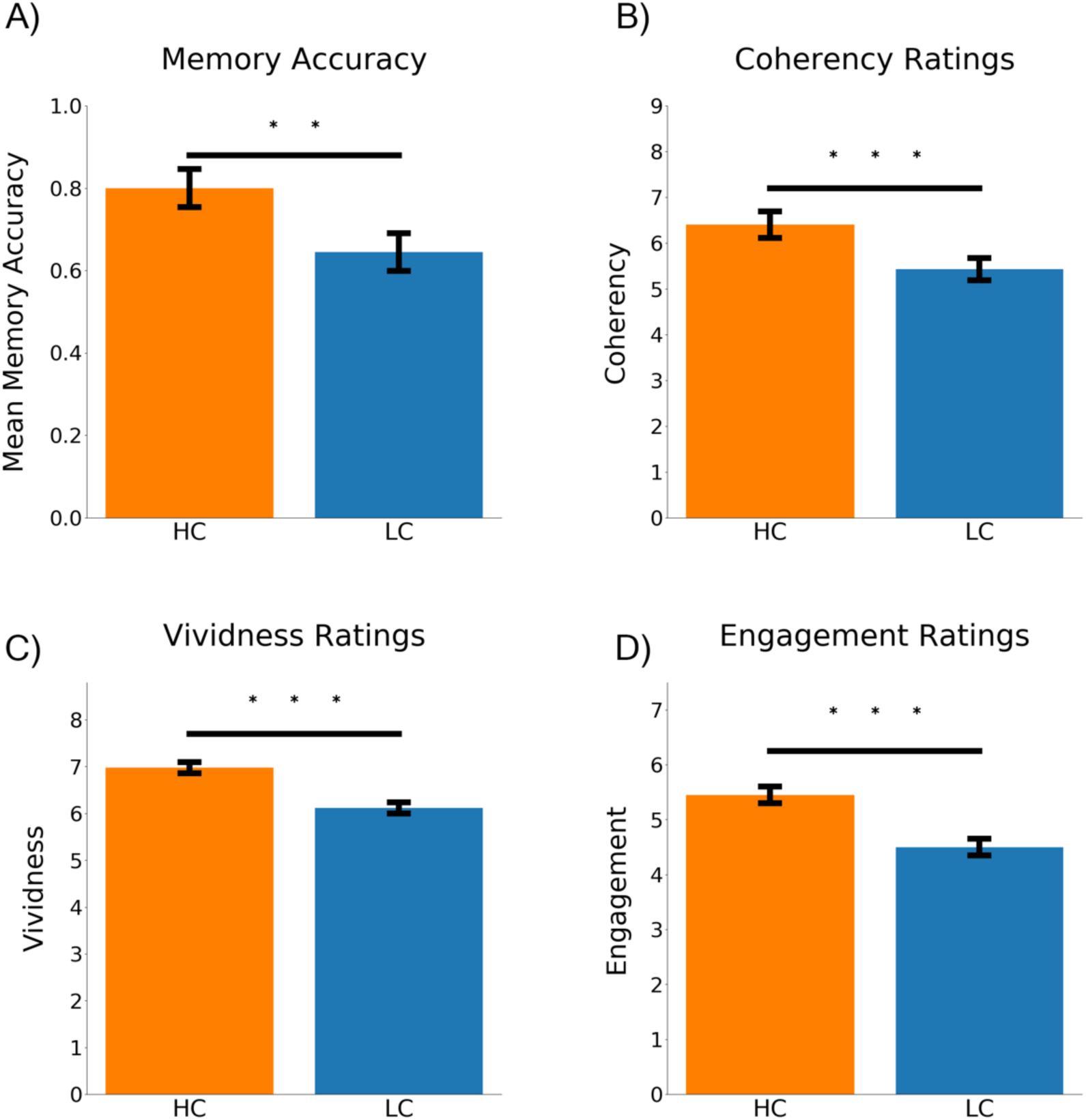
Behavioural results. A) Shows average memory performance for questions on HC versus LC videos. B) Shows that participants on average found HC videos more coherent. C) Higher vividness ratings were observed for HC videos. D) Participants reported HC videos as more engaging. ** p < .01; *** p < .001

The follow-up online study replicated all the behavioural results from the fMRI study (see Fig. 3). Participants freely recalled on average 76.8% of the video clips and performed better on the HC videos (84.35%) when compared to the LC videos (69.4%) (Z = 6.76 p < 0.001). Participants also rated their memory for the HC videos as more vivid (59.75) compared to the LC clips (51.04)(t_24.02_= 6.62, p < 0.001) and more coherent (HC, 66.33, LC, 54.78; t_65.61_ = 8.79, p < 0.001).

**Figure 3.**
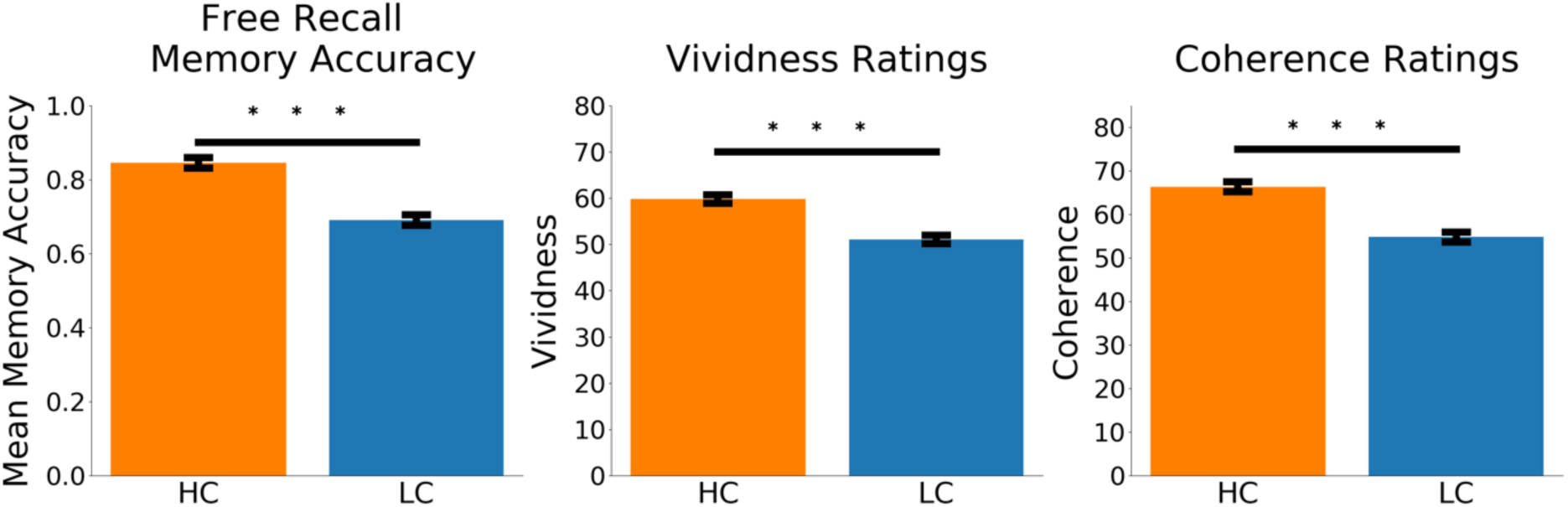
Results from Online Experiment. We show average free recall, vividness and coherence ratings for the HC and LC videos. *** p < 0.001.

The main purpose of the online study was to examine the semantic consistency across participants of their recalled descriptions of second-half clips. We also used Google’s Universal Sentence Encoder (USE) to quantify the semantic similarity of free recall responses for the videos in HC and LC conditions. Specifically, we examined how similar individual participants’ free recall responses were to the group average free recall responses, and, critically, whether the similarity was higher when the group had watched the video in the HC, compared to the LC condition. Consistent with the view that provision of prior knowledge guides and constrains how the content of the concluding part of the dialogue is interpreted and represented, we found that second-half descriptions were more similar across participants when the videos were seen in the HC condition when compared to the LC condition (t_19_ = 3.71, p = 0.001; See Supplementary Fig. 7).

#### fMRI Results

##### GLM

The contrast of watching videos versus the active baseline task revealed higher activation in visual, auditory, medial and anterior temporal cortices (see Supplementary Fig. 1). This is consistent with previous studies using videos (e.g. Bartels & Zeki, 2004). Contrasting first half clips with normal speech (NS) vs videos with spectrally rotated speech (SRS), identified regions commonly associated with language and semantic tasks (see Supplementary Fig. 2). Brain areas that showed higher activation for the onset of the videos versus baseline are shown in Supplementary Fig. 3. These results replicate Keidel et al.’s (2017) findings and indicate that retrosplenial cortex (RSC) and parahippocampal cortex (PHG) extending into the ventral precuneus showed transient activation associated with the onset of the videos. The results of the FIR time-course analysis were also consistent with this previous study in showing higher activation in the middle temporal gyrus (MTG), supramarginal gyrus (SMG) and angular gyrus (AG) for videos depicting a continuation of a previous narrative (HC clips). See Supplementary Fig. 4 and Materials for further discussion.

##### Inter-Subject Correlation

The ISC analysis allowed us to examine synchronization of the BOLD response across participants. ISCs for watching videos irrespective of the context manipulation were found in extensive regions of the occipital and superior temporal lobes, encompassing visual and auditory processing regions. Higher synchronization was also observed in bilateral IFG, medial prefrontal cortex, AG and precuneus (see Fig. 4).

**Figure 4.**
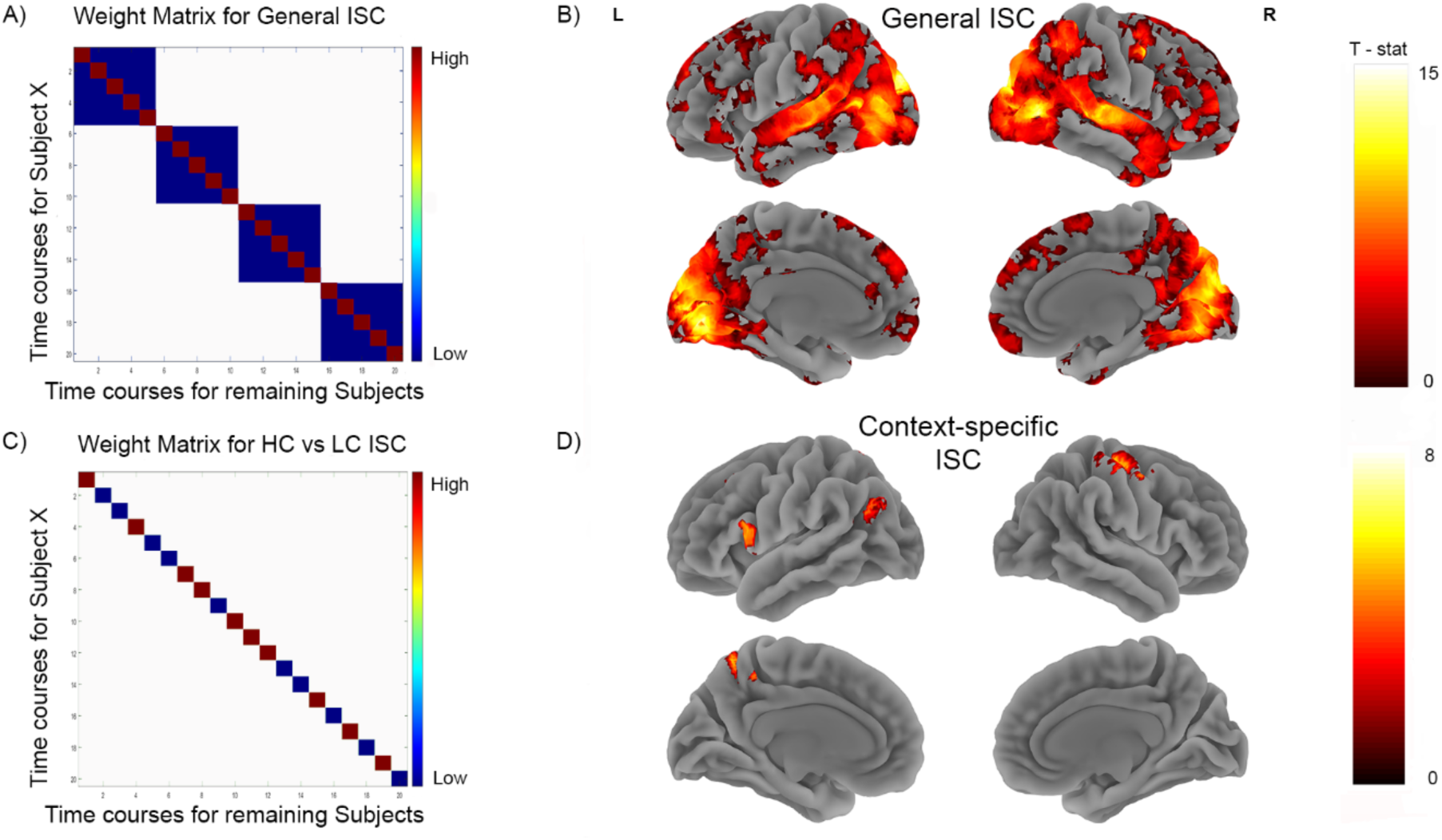
Inter-subject correlations. A) The weight matrix (General ISC) tests for video specific time course similarity across participants. Each cell represents the correlation between subjects’ time course for a particular video with the average time course of all remaining participants for a particular video. The diagonal represents correlations between time courses for the same videos. The off diagonal represents temporal correlations between mismatching videos within the same run. B) Brain map from video specific analysis, which shows extended synchronization across the brain for people watching the same videos. C) Weight matrix that tests for the time-course similarity across the same videos, depending on the prior knowledge provided for them. D) Brain map showing how time-course synchronicity was modulated by prior knowledge. Both brain maps show clusters significant at FWE p < 0.05 after permutation testing.

The key ISC comparison between HC and LC videos showed stronger coupling in the left AG, IFG, superior parietal lobule (SPL), superior frontal gyrus, and right precentral gyrus (see Fig. 4). In these regions, the synchrony across individuals is significantly greater when those individuals share knowledge of the preceding narrative.

##### Inter-Subject Pattern Similarity

The general ISPS allowed us to examine where spatial patterns of activity were more similar across participants watching the same videos versus watching different videos irrespective of the context manipulation. The results are shown in Fig. 5. This analysis revealed significant video-specific similarity across participants in the primary sensory areas and areas associated with higher cognitive processes, such as the bilateral IFG, precuneus, the MTG, the inferior portion of the anterior temporal pole (ATL) and the medial prefrontal cortex. The video-specific pattern similarity results reported here strongly resemble the event-specific intersubject similarity effects reported by Chen et al. (2017) and others.

**Figure 5.**
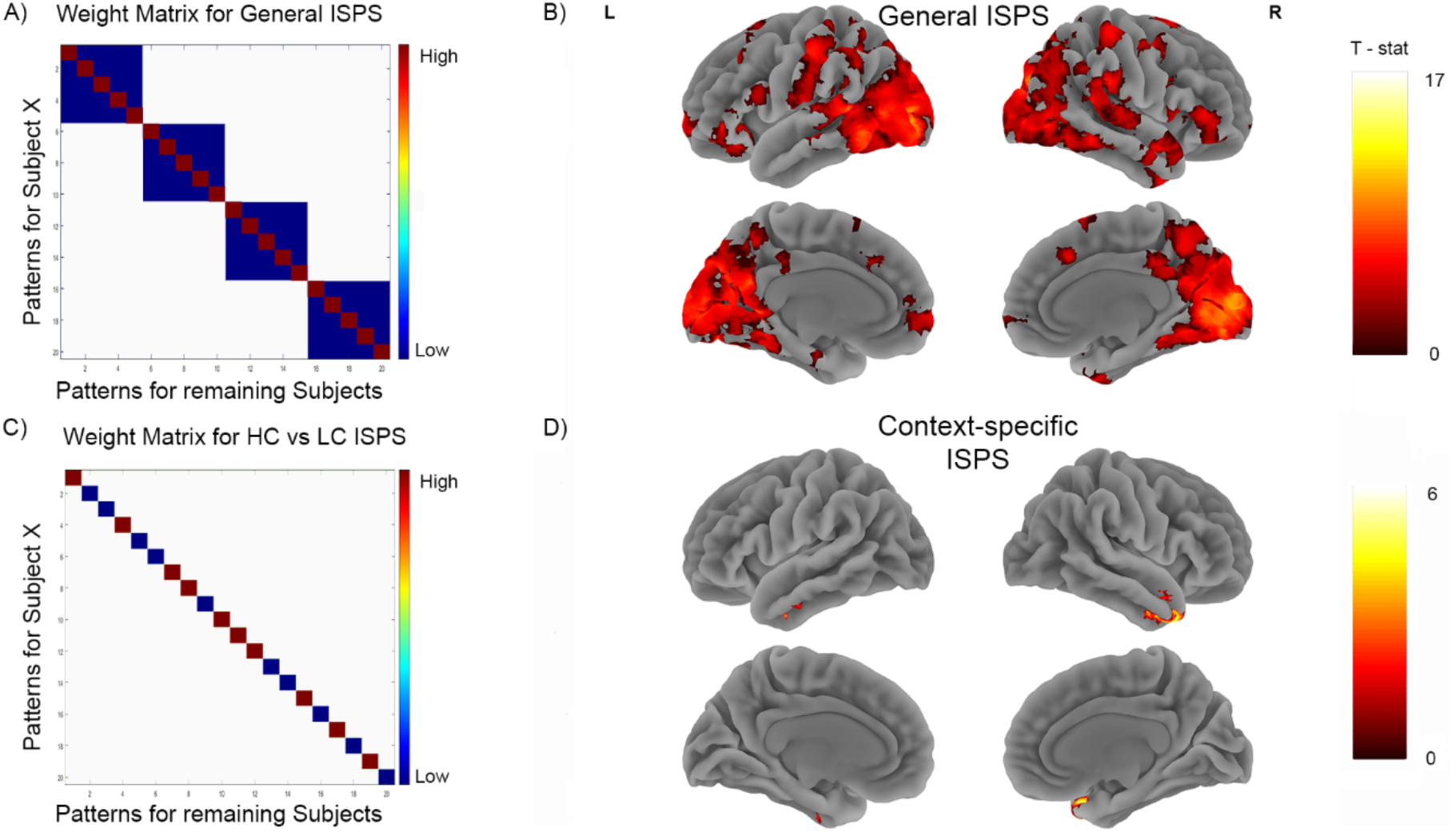
Inter-subject Pattern Similarity. A) The weight matrix (General ISPS) tests for video specific spatial pattern similarity across participants. The diagonal represents spatial similarity for the same videos. The off diagonal represents spatial correlations between mismatching videos within the same run. B) Brain map from video specific analysis, which shows extended pattern similarity across the brain for people watching the same videos. C) Weight matrix that tests for the spatial similarity across the same videos, depending on the prior knowledge provided for them. D) Brain map showing how spatial pattern similarity was modulated by prior knowledge. Both brain maps show clusters significant at FWE p < 0.05 after permutation testing.

However, our main interest was to examine whether narrative theme modulates this effect. The HC > LC contrast reveals effects in the left and right anterior temporal lobe (ATL). In these regions, the average pattern of activity across the whole second-half video clip was more similar between individuals who shared prior knowledge about the topic of the dialogue.

Recent studies have found some evidence that the hippocampus plays a role in integrating narrative information across delays or conflicting storylines (Chang et al., 2021; Cohn-Sheehy et al., 2020; Milivojevic et al., 2016). Accordingly, we also ran the ISPS HC > LC contrast within left and right hippocampal ROIs (see ‘*HC vs LC ISPS effects in hippocampus’* in supplementary materials). This post-hoc exploratory analysis revealed marginally significantly higher pattern synchronization in right hippocampus when participants were watching the HC clips compared to the LC clips (t_18_ = 2.09, p = 0.027, one-tailed). This is weakly supportive of a role for the hippocampus in retrieving and integrating the content of the first-half dialogue in order to comprehend and remember the second-half.

## Discussion

This study examined the cognitive and neural effects of prior knowledge of a narrative on the processing of dialogues between two people. Participants viewed videos of dialogues for which they either were, or were not, provided with knowledge of the preceding narrative (HC and LC respectively). Prior knowledge increased participants’ memory for the dialogues as well as their subjective comprehension and vividness ratings. A follow-up online study of 168 participants replicated these effects and found that freely recalled descriptions of the second-half dialogues were more semantically similar across participants in the HC condition. A number of brain regions showed higher ISCs in the HC condition, particularly those associated with the brain’s semantic network (including the left superior and inferior frontal gyrus, and left AG). In addition, we observed greater ISPS for HC videos in another region strongly associated with semantic knowledge, the bilateral anterior temporal lobes. Since the only difference between the two conditions was the provision of information about the topics of the dialogues, the fMRI effects likely represent the activation of this knowledge when participants watched the continuations of the dialogues. Therefore, our results highlight a role of the semantic network in integrating narrative information together in order to understand everyday conversations.

Prior knowledge boosted cued-recall memory accuracy for details from the second-half clips. It also boosted subjective ratings of how coherent and engaging the second-half clips were as well as how vividly they could be remembered. These effects are consistent with previous studies (Ames et al., 2015; Bellana et al., 2019; Bransford & Johnson, 1972; Brod et al., 2016; Dooling & Lachman, 1971; Johnson et al., 1974; Keidel et al., 2017; Liu et al., 2016; Raykov et al., 2020, 2021; St. George et al., 1994; van Kesteren et al., 2014). We also hypothesised that prior knowledge of the first-half of the dialogues would constrain how the second-half dialogues were interpreted, with the result that participants who shared knowledge of the first-half dialogues would process and remember the second-half dialogues in a more consistent manner. To test this hypothesis, we conducted a follow-up study that used a similar behavioural paradigm and was carried out online. After encoding the HC and LC clips participants’ memory for the second half clips was tested with a free recall test. A recently developed language model (Google USE; Cer et al., 2018) allowed us to quantify the similarity between free recall responses across participants. The freely recalled descriptions of the second-half clips were more similar across participants who had watched the first-half clips with intelligible speech.

To illustrate this effect with an example; in one of the second-half videos, a character claims that penguins once “filled the skies” but are now unable to fly because they lack the confidence to do so. A participant viewing this clip in the HC condition can relate this discussion back to the first-half, in which the characters discuss how low self-esteem had a negative effect on a colleague’s work. However, participants in the LC condition did not have this background context to draw on and tended to form more idiosyncratic interpretations of the scene. Our fMRI findings demonstrate that sharing knowledge of the preceding dialogue not only results in greater alignment in the interpretation of the second half clips but also greater alignment in brain activity across individuals. This provides strong evidence that the brain regions we identify play a role in integrating narrative information across delays in order to understand conversations.

The brain regions where activity was modulated by prior knowledge are a subset of those that show more general memory reinstatement effects (Bird et al., 2015; Chen et al., 2017; Oedekoven et al., 2017; Polyn et al., 2005; Raykov et al., 2021; St-Laurent et al., 2015; Zadbood et al., 2017), and they have been particularly associated with “semantic control” (Binder et al., 2009; Lambon Ralph et al., 2016; Noonan et al., 2013). The MTG, IFG and AG were recently highlighted in a similar fMRI study involving sitcoms (Keidel et al., 2017). Activity in these regions was higher when a background narrative context was provided. The authors suggested that these regions processed the “on-the-fly” links between semantic concepts necessary to comprehend the storyline. In this study, we carried out the same analysis as Keidel et al., (2017) and replicated their findings for the MTG and AG, but not the IFG (see Supplementary Fig. 4). Furthermore, a number of recent studies using spoken short stories have also implicated the IFG, AG and lateral temporal regions in the integration of prior knowledge with incoming information in order to understand the unfolding narrative (Chang et al., 2021; Yeshurun et al., 2017). More generally, when people interpret a narrative in a similar way, brain activity is more synchronised within these regions (Nguyen et al., 2019 report effects in DMN regions; Saalasti et al., 2019 see syncoronization in AG). Causal evidence for a role of the AG in the integration of prior information to understand a narrative comes from a recent study by Branzi and colleagues (2021) who used TMS on the left AG which resulted in disrupted context-dependent comprehension of prose passages. Taken together, these studies suggest that the MTG, IFG and AG support the activation, integration and representation of conceptual information required to understand dialogues as they unfold over time.

These empirical findings mesh well with recent theoretical accounts of processing with DMN structures. The inferior lateral parietal cortex has been viewed as supporting a “buffer” for episodic information (Baddeley, 2000; Vilberg & Rugg, 2007), or for linking together the features of an event into a cortical representation (Ramanan et al., 2018; Shimamura, 2011). Others have argued that its’ role is in supporting abstract conceptual representations of events (such as a “birthday party”, Binder & Desai, 2011). More recent views have downplayed the traditional dichotomy of episodic versus semantic memory (Renoult et al., 2019). For example, Humphreys et al., (2021) argued that the angular gyrus operates as a multisensory buffer, supporting “consciously accessible representations that integrate features of events that unfold over time” (Hasson et al., 2015; see also Hasson et al., 2008; Lerner et al., 2011).

We additionally carried out ISPS analyses to see where the spatial patterns of BOLD activity were more consistent between participants sharing narrative contextual information. ISPS may provide complimentary insights compared to ISC analyses (Nastase et al., 2019). These effects were localised to the ATL bilaterally. Although these results are due to the same contrast as the ISC results (greater similarity for the HC compared to the LC clips), they may be driven by subtly different mechanisms. While ISC effects reflect temporal variations in activity during the clips that are more synchronised across participants, the ISPS effects are driven by spatial variation in the activity patterns for each clip, where the patterns reflect time-averaged responses. Therefore, these effects might represent shared representations of the overarching narrative themes (Chen et al., 2017), rather than more dynamic processing of concepts relating to the ongoing narrative. The importance of the ATL for semantic memory and conceptual knowledge is well-established (L. Chen et al., 2017; Patterson et al., 2007; Rice et al., 2015). Within the ATL, the anterior and ventral regions identified in our study have been argued to act as a semantic “hub”, supporting amodal conceptual representations that are independent of specific sensory input (Murphy et al., 2017; Patterson et al., 2007). Our findings suggest that the ATL support abstract representations of events, perhaps similar to the “event semantics” described by Binder and Desai (2011).

Many studies have highlighted a particular role for the hippocampus in the integration of information held in memory (Backus et al., 2016; Clewett et al., 2019; Cohn-Sheehy et al., 2020; Griffiths & Fuentemilla, 2019; Schlichting & Preston, 2015, 2016; Shohamy & Wagner, 2008; Zeithamova & Bowman, 2020; Zeithamova, Dominick, et al., 2012; Zeithamova & Preston, 2010; Zeithamova, Schlichting, & Preston, 2012). We therefore carried out an exploratory analysis to examine whether spatial patterns of BOLD activity were more similar within the hippocampus across individuals for the HC videos. We did indeed find significantly higher ISPS for the HC videos in the right hippocampus at a marginally significant level. Interestingly, two previous studies that implicated the hippocampus in the integration and differentiation of narrative storylines also found effects in the right, but not the left, hippocampus (Cohn-Sheehy et al., 2020; Milivojevic et al., 2016). These findings support the view that the hippocampus plays a role in activating and integrating information from memory in order to process current experiences.

Of course, we would expect multimodal events to be represented across many brain regions. If we take together these previous findings and the results of the current study, it appears that there is a division of labour in the instantiation of event models in the brain. Regions of the DMN, notably posterior midline regions, represent the stable core of the model, such as the location and identity of the people present (Chen et al., 2017; Robin et al., 2018). By contrast, the ATL may play a central role in representing the overarching narrative gist. Finally, other regions of the semantic network, particularly the IFG and AG, may more tightly track the changing aspects of the associated narrative and link them to prior knowledge (see also Ranganath & Ritchey, 2012; Ritchey & Cooper, 2020).

In summary, knowledge about a preceding dialogue enabled participants to better comprehend and remember the conclusions of these dialogues. The availability of narrative information led to increases in intersubject fMRI synchronization and spatial pattern similarity, predominately in regions associated with semantic processing. Our findings highlight a role for these regions in the retrieval and activation of concepts necessary to understand a complex dialogue, since other aspects of the events, such as the familiarity of the people and locations were held constant. These results provide important new insights into how the brain represents and updates narrative information as well as highlighting an important case of functional specialisation within the wider network of brain regions that are involved with event processing.

## Acknowledgements

This work was supported by the European Research Council (337822 to C.M.B.) and an Economic and Social Research Council studentship (ES/J500173/1 to P.P.R.) and fellowship (ES/V012444/1 to P.P.R.).

