## Supplementary Materials for "The importance of semantic network brain regions in integrating prior knowledge with an ongoing dialogue"

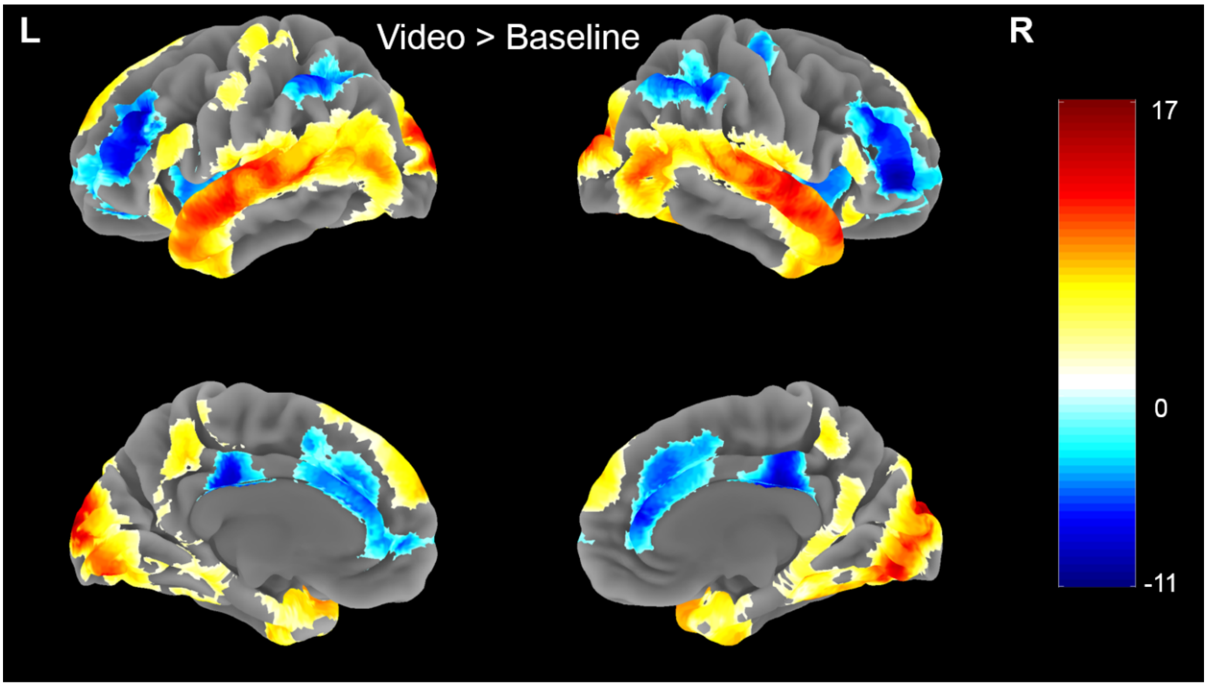


**Supplementary Figure 1**: *Video vs Baseline analysis.* The map shows brain regions active throughout the duration of the videos (from all conditions) versus baseline.

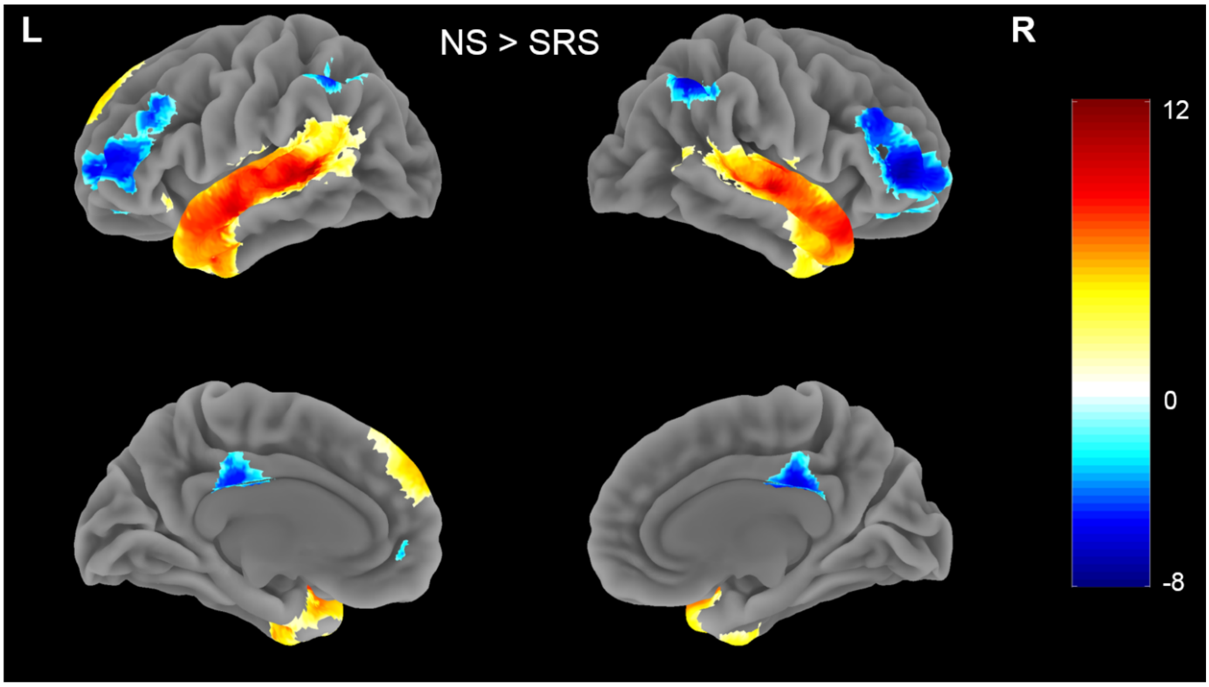


**Supplementary Figure 2**: *Language contrast.* Brain map showing univariate contrast of first half videos with intact speech (NS) versus first half videos without comprehensible speech (SRS).

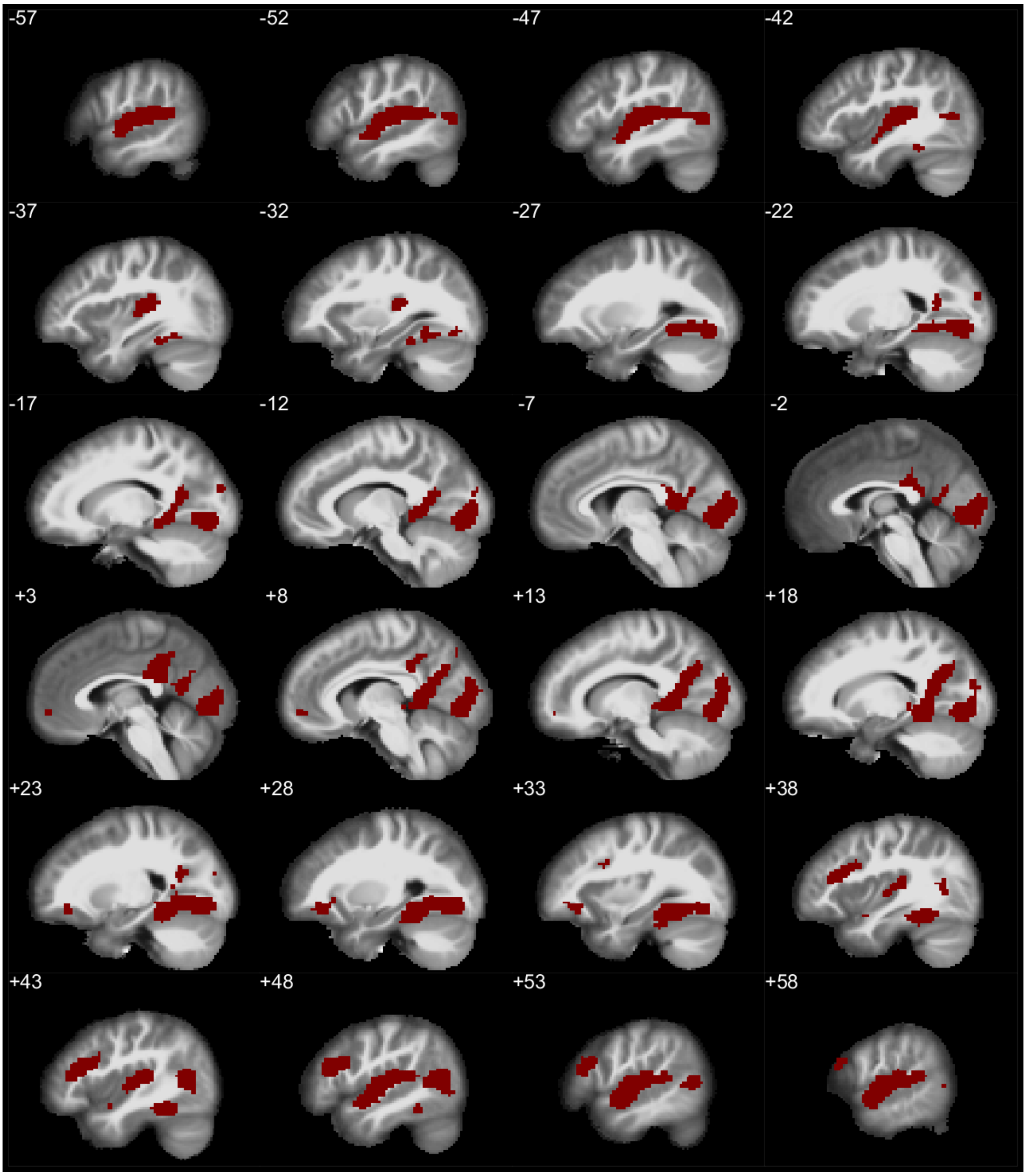


**Supplementary Figure 3**: *Analysis of video onsets.* Conjunction analysis showing the positive onset responses for HC, LC and NS videos. Voxels in red represent whole-brain significant ($p < 0.001$) responses to all 3 conditions.

### Supplementary Methods

We implemented a FIR time-course analysis in order to analyse how activity differed over the duration of the videos. A set of 12 piecewise linear tent functions were used to model the first 26.2 seconds of each of the 4 conditions. Keidel et al. (2017) found significantly greater activation in MTG, AG, SMG and IFG for HC as compared to LC videos. To attempt to replicate this finding we averaged the results of the time course analysis in these clusters. Significant differences in the time-course were found in the left MTG, AG, and supramarginal gyrus (SMG) replicating the previous findings, while no significant difference was observed in the left IFG.

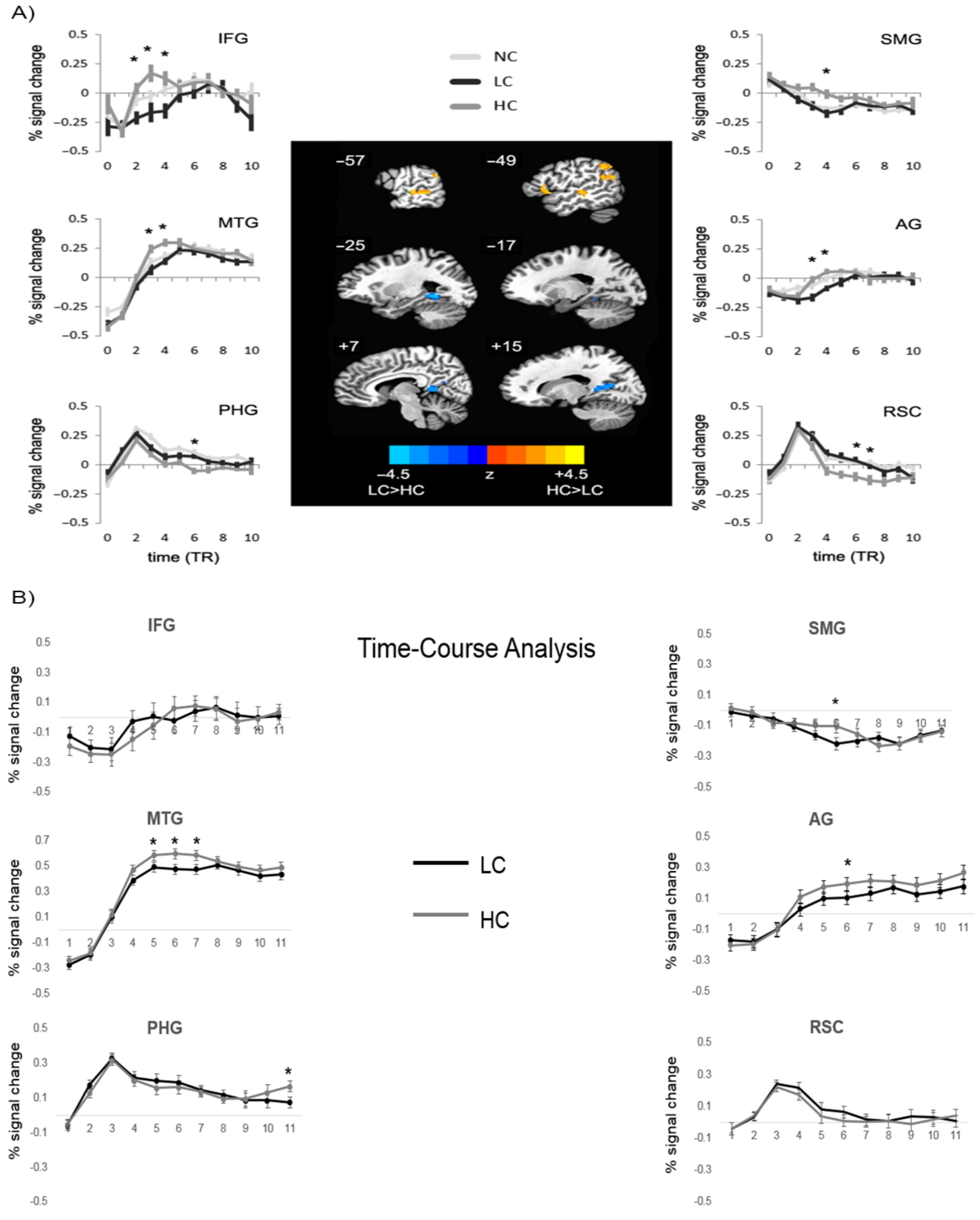


**Supplementary Figure 4**: Deconvolution FIR analysis examining response to the initial 26.2 seconds for HC and LC videos. A) Results reported by Keidel et al. (2017). B) Time courses in ROIs identified in Keidel et al. (2017) for HC versus LC videos.

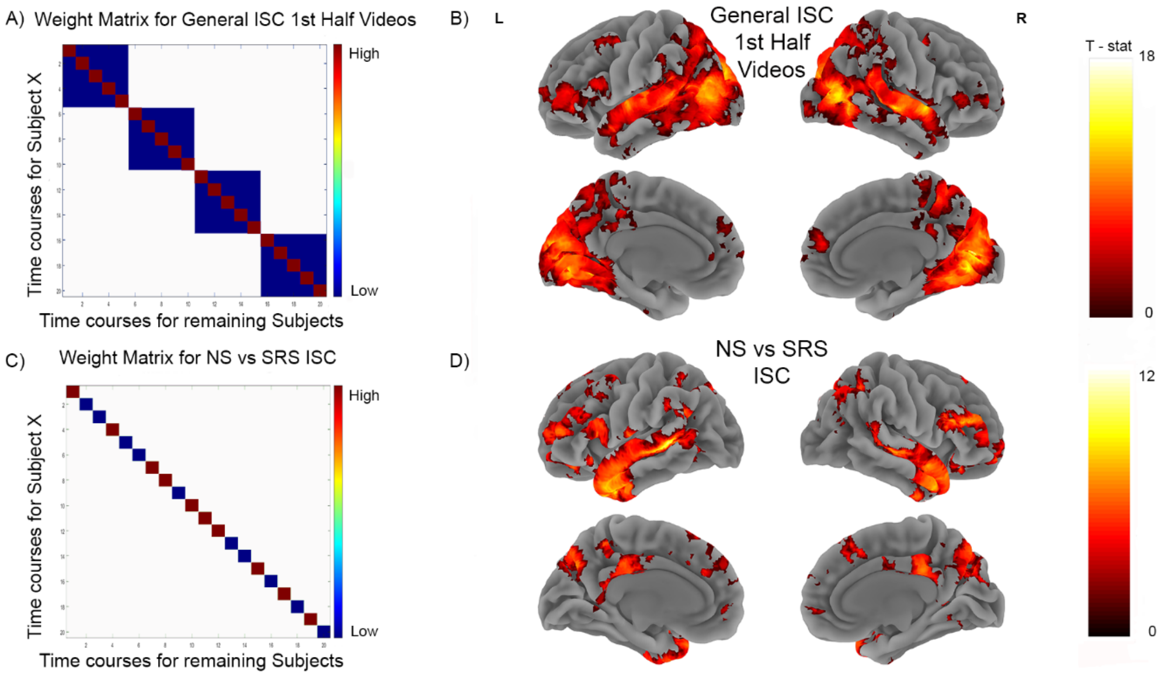


**Supplementary Figure 5:** *Inter-subject correlations for first half videos*. A) The weight matrix (General ISC) tests for video specific time course similarity across participants. Each cell represents the correlation between subjects' time course for a particular video with the average time course of all remaining participants for a particular video. The diagonal represents correlations between time courses for the same videos. The off diagonal represents temporal correlations between mismatching videos within the same run. B) Brain map from video specific analysis, which shows extended synchronization across the brain for people watching the same videos. C) Weight matrix that tests for the time-course similarity across the same videos, depending on whether the language in the videos is intelligible (NS) or not (SRS). D) Brain map showing how time-course synchronicity was modulated by provision of comprehensible language in the videos. Both brain maps show clusters significant at FWE $p < 0.05$ after permutation testing.

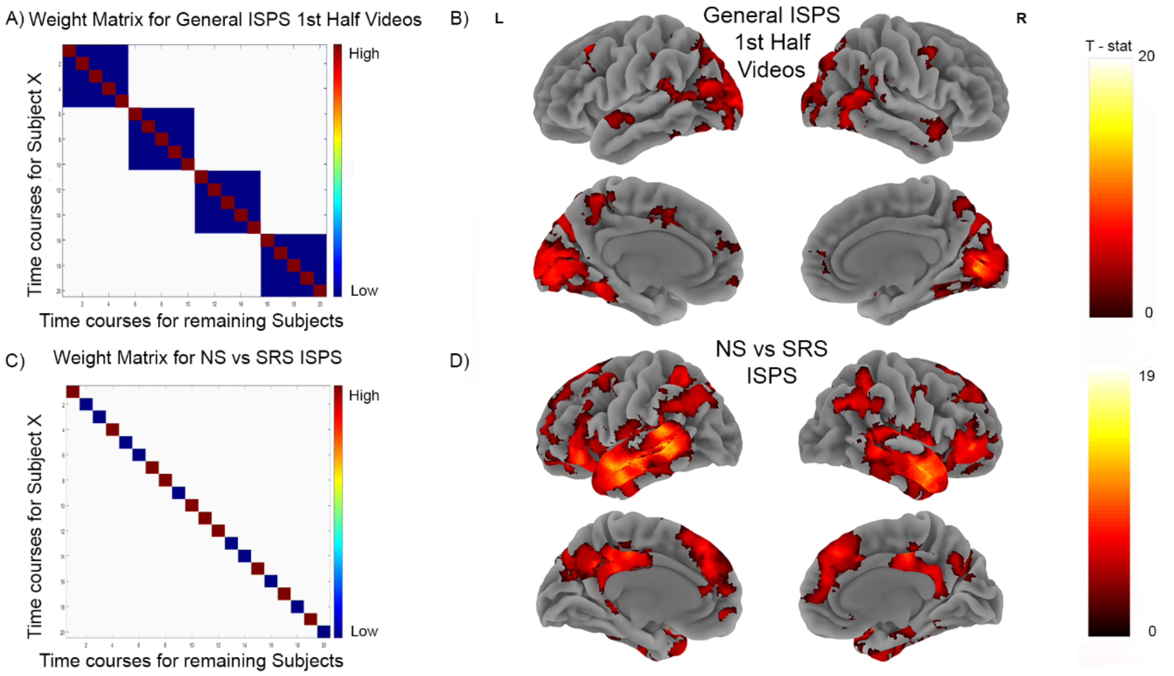


**Supplementary Figure 6**: *Inter-subject Pattern Similarity for first half videos*. A) The weight matrix (General ISPS) tests for video specific spatial pattern similarity across participants. The diagonal represents spatial similarity for the same videos. The off diagonal represents spatial correlations between mismatching videos within the same run. B) Brain map from video specific analysis, which shows extended pattern similarity across the brain for people watching the same videos. C) Weight matrix that tests for the spatial similarity across the same videos, depending on the whether the language is comprehensible (NS) or not SRS. D) Brain map showing how spatial pattern similarity was modulated by presence of comprehensible narrative. Both brain maps show clusters significant at FWE $p < 0.05$ after permutation testing.

#### HC vs LC ISPS effects in Hippocampus

A recent study examined whether patterns of activity in hippocampus were modulated when participants could integrate narrative events that were presented far apart. Participants listened to short stories in an interleaved manner. Some of the stories were related and participants could integrate over them to form a coherent narrative, other stories could not be integrated into coherent narratives. Cohn-Sheehy and colleagues (2020) found that patterns of activity in right hippocampus differentiated between the stories that formed coherent narratives and the stories that did not. Patterns of activity in right hippocampus were more similar when participants were listening stories that could be integrated when compared to when they were listening to stories that could not be integrated. Other studies using long interleaved narratives have also observed that hippocampus plays an important role in differentiating between the stories (Chang et al., 2021; Milivojevic et al., 2016; Milivojevic, Vicente-Grabovetsky, & Doeller, 2015). Based on this prior work we ran analysis examining whether ISPS effects would be higher when participants were watching the HC videos when compared to the LC videos. Participants had knowledge of the narrative preceding the HC video and therefore could integrate the newly incoming information with their prior knowledge of the dialogue, which they could not do for the LC videos. We examined ISPS effects in left and right hippocampus. Both hippocampal ROIs were based on from MTL segmentation protocol outlined in (Ritchey, Montchal, Yonelinas, & Ranganath, 2015) (see https://neurovault.org/collections/3731/). We combined the head, body and tail of hippocampus to form a single ROI for each hemisphere. ISPS effects did not differ between the HC and LC conditions in the left hippocampus ($t_{18} = 0.77$, $p = 0.45$). However, right hippocampal ROI showed higher synchronization across participants when participants were watching the HC videos when compared to the LC videos ($t_{18} = 2.09$, $p = 0.027$ – one tailed). These results are in line with results from Cohn-Sheehy (2020) and other work showing that hippocampus plays an important role in integration (Schlichting & Preston, 2015; Shohamy & Wagner, 2008; Zeithamova & Bowman, 2020; Zeithamova, Dominick, et al., 2012). However, we note that our hippocampal analyses were exploratory and should be interpreted with caution.

**Semantic Consistency in Free Recall responses**

The participants in the online study provided free recall responses for each of the video clips. The responses were scored with a binary Remember/No score. Apart from investigating differences in memory accuracy we also examined whether there was higher semantic consistency between free recall responses when participants watched the video under the HC condition compared to the LC condition. Specifically, we used recently developed deep neural network that allows embedding of text data into high-dimensional vectors. Importantly, these high-dimensional vectors are constructed in such a way that they maintain the semantic similarity between sentences. We used the Google's Universal Sentence Encoder (USE), which allowed us to quantitively examine the similarity between free-recall responses across participants. Specifically, we focused only on trials that were correctly remembered. We transformed the correctly remembered free recall responses into high-dimensional semantic vectors using USE. Afterwards, for each video we examined how similar were the participants responses were to other participant's responses that watched the video under the same condition. For example, for video one, 74 participants correctly remembered it and had seen it as HC video, and another 71 participants correctly remembered it and had seen it as LC video. We then constructed two average semantic vectors for video 1. One was the average semantic vector of everyone that remembered the video in the HC condition, and the other was the mean semantic vector for everyone that remembered the video in the LC condition. We computed the similarity between each participant's free recall response to the average semantic vector for that video watched under the same condition. This allowed us to examine the semantic consistency or spread of responses in semantic space for each video under each condition. We used Pearson's correlation coefficient to compute the similarity between the embedded recall descriptions. Note the average semantic vectors were always recalculated to not include the description of the current participant (using a "leave one subject out" procedure). After computing each participants similarity to the average semantic pattern we Fisher transformed and averaged these similarity scores to compute an average semantic consistency measure for each video both in the HC and LC condition. We then computed a non-parametric permutation tests to examine whether semantic consistency was higher for videos watched under the HC condition than the LC condition (see Supplementary Fig. 7). This analysis and its results are also reported in the main text.

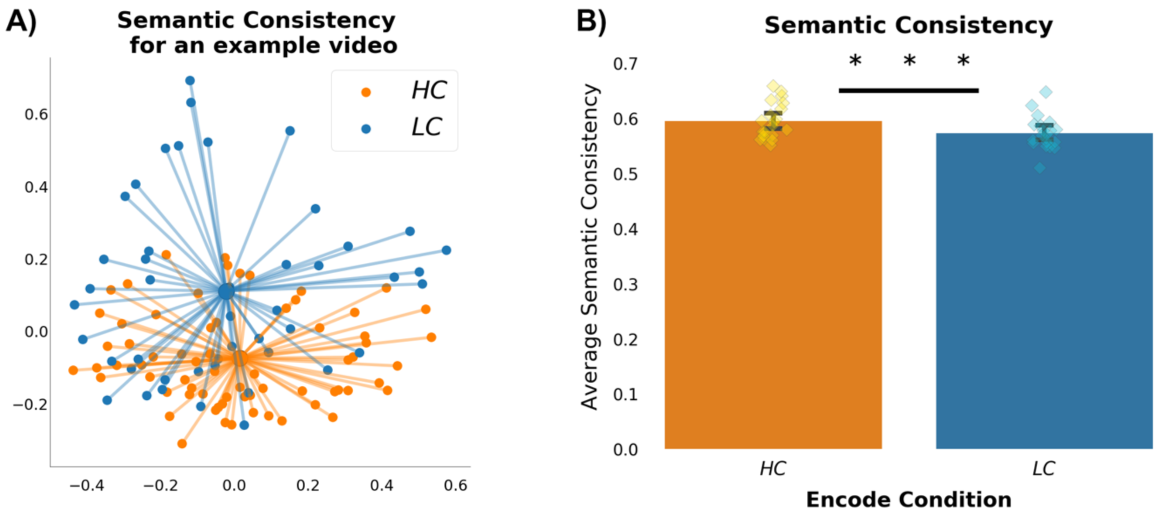


**Supplementary Figure 7:** Semantic Consistency. A) Shows a PCA projection in 2 dimensions of the semantic vectors that represent participants correct free recall responses for a single video. Semantic consistency for each video was calculated within each condition. The larger centre dot is the average semantic pattern for the video based on the free recall responses. Lines towards the centre are for illustrative purposes. B) We calculated the average spread of scores for each video under each condition. One data point in the bar graph represented the average spread of responses across participants watching a video in one condition (e.g. average spread in orange points in A)). Semantic consistency was on average higher for the HC videos. * * * - p = 0.001
